# Finding and Keeping the Beat: Neural Mechanisms Differ as Beat Perception Unfolds

**DOI:** 10.1101/2021.10.01.462808

**Authors:** Daniel J. Cameron, Jessica A. Grahn

## Abstract

Perception of a regular beat is essential to our ability to synchronize movements to music in an anticipatory fashion. Beat perception requires multiple, distinct neural functions, corresponding to the perceptual stages that occur over time, including 1) detection that regularity is present (*beat finding*), 2) prediction of future regular events to enable anticipation (*beat continuation*), and 3) dynamic adjustment of predictions as the rhythmic stimulus changes (*beat adjustment*). The striatum has been shown to be crucial for beat perception generally, although it is unclear how, or whether, distinct regions of the striatum contribute to these different stages of beat perception. Here, we used fMRI to investigate the activity of striatal subregions during the different stages of beat perception. Participants listened to pairs of rhythms (polyrhythms) whose temporal structure induced distinct perceptual stages—*finding, continuation*, and *adjustment* of the beat. Dorsal putamen was preferentially active during beat *finding*, whereas the ventral putamen was preferentially active during beat *adjustment*. We also observed that anterior insula activity was sensitive to metrical structure (greater when polyrhythms were metrically incongruent than when they were congruent). These data implicate the dorsal putamen in the detection of regularity, possibly by detection of coincidences between cortical oscillations, and the ventral putamen in the adjustment of regularity perception, possibly by integration of prediction errors in ongoing beat predictions. Additionally, activity in the supramarginal and superior temporal gyri correlated with beat tapping performance, and activity in the superior temporal gyrus correlated with beat perception (performance on the Beat Alignment Test).

## Introduction

Detecting and anticipating regular events in the environment are critical functions of the human brain. Sensitivity to certain auditory temporal regularities allows the detection and anticipation of beats (the regular emphases sensed in musical rhythms). Beat perception enables humans to accurately synchronize movement to rhythm (e.g., tapping a toe on, not after, the beat; Aschersleben, 2002), a behavior that is not generally observed in non-human primates (Zarco, Merchant, Prado, & Mendez, 2009). Beat-synchronized movements (e.g., tapping) are flexible and can adapt to and persist through complex rhythms with distinct stages (Cameron & Grahn, 2014). The striatum is heavily implicated in beat perception (Grahn & Brett, 2007, 2009; Grahn & Rowe, 2009, 2013; Kung, Chen, Zatorre, & Penhune, 2013; Teki, Grube, Kumar, & Griffiths, 2011), although different areas (putamen, pallidum, and caudate) have been reported in different studies.

Variation in the location of reported striatal activity during previous studies may be because several different striatal functions mediate different stages of beat perception. For example, on first hearing a rhythm, regularity must be detected *(beat finding)*. One possible mechanism of *beat finding* is that medium spiny neurons in the striatum detect the coincidence of intrinsic neural oscillations with neural oscillations driven by rhythmic sounds (Matell & Meck, 2004). The detection of these coincidences by the striatum on a regular or predictable basis may result in entrainment of oscillations in neural systems (including motor and attentional systems) to the regularities in auditory rhythms. The initiation of entrainment may correspond to the onset of beat perception, or *beat finding*. A striatal role of detecting auditory regularity (the beat) in musical rhythms via coinciding cortical oscillations is consistent with prominent ‘neural resonance’ theories that beat perception arises from the interactions between banks of neural oscillators that entrain to auditory rhythms (Large & Kolen, 1994; Large & Snyder, 2009, Tal, Large, Rabinovitch, Schroeder, Poeppel, & Golumbic, 2017).

A second function that may be important for beat perception is temporal prediction. Temporal prediction is a hallmark of beat perception (humans tend to tap slightly before the beat (Aschersleben, 2002), and the striatum is known to support the generation of temporal predictions based on regularities in auditory stimuli (Kotz, Schwartze, & Schmidt-Kassow, 2009). Temporal predictions result in the beat percept persisting over time, even through silence or parts of a rhythm where the beat may be ambiguous, and thus support ongoing beat perception (*beat continuation*).

A related, third, striatal function may be processing of temporal prediction *errors* (McClure, Berns, & Montague, 2003), critical because beat perception unfolds over time and requires ongoing integration of new temporal information to update the predictions of future beat positions. This integration could support the persistence of beat perception through the speeding and slowing of music, or through polyrhythmic music that features conflicting beat structures: unexpected timing of incoming intervals is integrated into ongoing temporal predictions associated with the perceived beat. The striatum may be key to integrating these temporal prediction errors to maintain a unified beat percept over time (*beat adjustment*).

Previous work supports that different regions of the striatum may be important at each stage: the caudate nucleus is preferentially active during *beat finding* (Kung et al., 2013), and the putamen during *beat continuation* (Grahn & Rowe, 2013). Both caudate and putamen respond to short beat-based rhythms, when *beat finding* is likely occurring (Grahn & Brett, 2007). Thus, the striatum has been implicated in both beat finding and continuation. However, no studies have directly examined whether distinct regions of the striatum respond during *finding, continuation*, and *adjustment* stages of beat perception. Therefore, we used fMRI to measure striatal activity during these three stages, which differentially require detection of regularity, generation of predictions, and integration of prediction errors, and compared activity in distinct regions of the striatum.

## Methods

### Participants

18 participants (8 male, mean age 25.4 years) provided written, informed consent before completing two behavioral tasks and undergoing fMRI scanning. All participants completed the musical training subscale of the Goldsmiths Musical Sophistication Index (GMSI; Müllensiefen, Gingras, Musil, & Stewart, 2014), as well as the Beat Alignment Test (BAT) from the GMSI. Participants had an average overall GMSI training score of 29.5 out of a maximum possible score of 49.

### Stimuli

Stimuli were pairs of rhythms presented together, with one rhythm composed of 375 Hz sine tones and the other composed of 500 Hz sine tones. All tones were 100ms, with a linear rise and fall over the first and last 50ms, respectively. Within individual rhythms, inter-onset intervals (IOIs) were durations of 1, 2, 3, or 4 units, in which 1 was equal to one of five absolute durations (180, 195, 210, 225, or 240ms), and the other units scaled proportionately by 2, 3, or 4. The rhythms in each pair were always based on the same absolute unit duration. Intervals within each rhythm were ordered to conform to either a duple or a triple meter (with a strong beat occurring on either every fourth unit or every third unit, respectively). Individual rhythms were composed of repetitions of basic patterns of intervals summing to 16 units for duple rhythms or 12 units for triple rhythms.

There were four trial types, based on whether the rhythm pairs began *simultaneously* or in a *staggered* fashion, and whether the pairs were metrically *congruent* (both duple or both triple) or *incongruent* (one duple and one triple) as shown in Figure 1. In *simultaneous* trials, the two rhythms began and ended together, whereas in *staggered* trials, one rhythm started and the other rhythm began after a duration equal to the duration of one repetition of the second rhythm. The intensity of the first audible repetition of the second rhythm increased linearly from silence (i.e., faded in).

**Figure 1.**
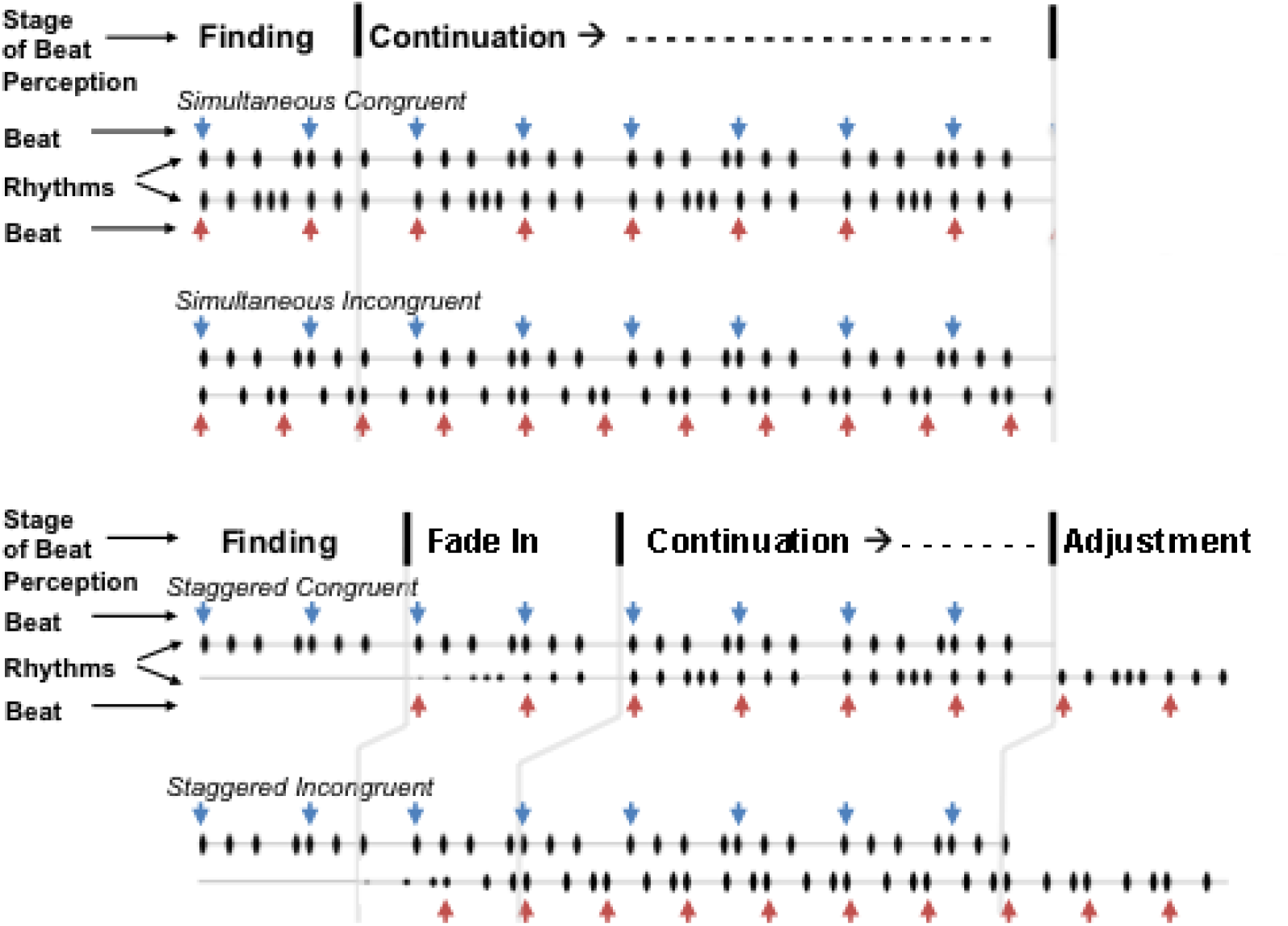
Examples of the four types of stimuli, each consisting of two rhythms. Waveforms of rhythms are presented in black. Blue and red arrows indicate beat positions. Rhythms were presented in simultaneous or staggered fashion. In simultaneous trials, the two rhythms began and ended together. In staggered trials, the second rhythm faded in starting after one cycle of the first rhythm (the fade-in lasted for one cycle of the second rhythm), and, after four cycles of the first rhythm, the first rhythm ended, as the second rhythm continued for one cycle. In congruent trials, both rhythms had the same beat rate (duple or triple) and in incongruent trials, one rhythm had a duple beat rate and the other had a triple beat rate. In the incongruent trials shown, the bottom rhythm has a triple meter and the top rhythm has a duple meter.

As behavioural evidence indicates only a singular beat percept emerges, even when multiple different simultaneous beat percepts are possible (Poudrier & Repp, 2013), and that people can maintain beat perception through *finding, continuation*, and *adjustment* (or switching) stages of *incongruent*, paired rhythms similar to those used here (Cameron & Grahn, 2014), we used metrically *incongruent* and congruent rhythm pairs to assess the neural mechanisms of beat perception in the context of one or multiple possible beat percepts. When multiple beat percepts are possible (i.e., metrical ambiguity, see London, 2004), additional neural mechanisms may be required to process rhythm and beat. For example, beat perception in metrically incongruent contexts may require weighting attention to the specific rhythm that corresponds to a particular beat, in order to establish a stable beat percept. Or, internal generation of temporal predictions may require greater neural resources in these contexts to overcome the conflicting information regarding beat percept.

### Behavioural tasks

Before scanning, participants completed a beat tapping task and a beat-strength rating task, using all stimuli used in the fMRI experiment. For each trial, participants tapped their perceived beat along with each rhythm. After each trial, they rated how difficult it was to maintain their beat percept on a 5-point scale. Stimuli were presented by laptop via noise-cancelling headphones outside of the scanner, and tapping and ratings were collected by laptop.

In the scanner, the first seven participants performed the same beat-strength rating task that they had completed before scanning (with no simultaneous beat tapping task). The subsequent 11 participants performed the same beat-strength rating task for 2 sessions and performed a deviant detection task in the other 2 sessions. For the deviant detection task, 14 of the 104 trials contained a deviant (square) tone in place of one of the regular tones. The deviant never occurred during the finding stage of a trial. For the 11 participants who completed both tasks, the tasks alternated over the four sessions in counterbalanced order over participants. For both beat rating and deviant detection tasks, responses were made on a 5-button response box. The addition of the deviant detection task was motivated by substantial activity in visual cortex observed in preliminary analyses of the first seven participants, thought to be due to the visual instruction screen presented before and after each trial (always immediately following the final stimulus presentation stage; the *continuation* stage in *simultaneous* trials, and the adjustment stage in *staggered* trials). The deviant detection task had no visual instruction after trials (participants were instructed in advance to respond if and when they heard a deviant), but required attending to the auditory stimuli (91.8% of trials were detected). For both tasks, 0, 4, 6, or 12 seconds of silence separated trials (evenly distributed except for the 0s silence, which occurred once per session). Stimuli were presented in the scanner via pneumatic Sensimetrics insert earphones.

### Image acquisition and fMRI design

FMRI data were collected on a 3-T Siemens Magnetom Prisma MRI scanner in 4 sessions of 240 echo-planar imaging (EPI) volumes. Each EPI volume had 43 slices of 3mm thickness, an interslice gap of 1mm, and a resolution of 3×3mm. Repetition time (TR) was 2.5s. All analyses were completed with SPM8 (SPM8; Wellcome Department of Imaging Neuroscience, London, UK). Anatomical images (magnetization prepared rapid gradient echo, or MP-RAGE) were collected for co-registration. For each participant, images were interpolated in time and spatially realigned to the mean image using 2^nd^ degree B-spline interpolation. The co-registered structural image was segmented using affine regularization and normalized at a resolution of 1×1×1 mm to a standard ICBM template in Montreal Neurological Institute space. EPI images were normalized to the template at 3×3×3 mm resolution and spatially smoothed with a 8mm full-width half-maximum Gaussian kernel.

Within-subject first level modeling included 14 conditions: Conditions 1 and 2) the *finding* and *continuation* stages in simultaneous incongruent trials, 3 and 4) the *finding* and *continuation* stage in simultaneous congruent trials, 5) the *finding* stage in staggered trials, 6-8) the *fade-in, continuation, and adjustment* stages in staggered incongruent trials, 9-11) the *fade-in, continuation, and adjustment* stages in staggered congruent trials, and (12-14) instruction screen viewing, responses, and deviant tones). The durations of rhythm conditions are depicted in Figure 1. For trials with simultaneously presented rhythms, *finding* epochs had a duration equal to one cycle of triple meter ranging from 2.0 – 3.8 s. *Continuation* epochs in simultaneous trials began immediately following the *finding* epoch and continued for the remainder of the stimulus (4 cycles of a triple meter rhythm or 3.25 cycles of a duple rhythm). For trials with staggered rhythms, the duration of *finding* epochs was equal to one cycle of the second rhythm (the time from stimulus onset to onset of the second rhythm). *Fade-in* epochs began at the onset of the second rhythm with the duration equal to one cycle of that rhythm (over which time the rhythm linearly increased from silent to full intensity). *Continuation* epochs began after the first cycle of the first rhythm (immediately following the *fade-in* epoch), and the duration was equal to 4 cycles if it was a triple meter rhythm or 3 cycles of the second rhythm if it was a duple meter rhythm. *Adjustment* epochs began immediately after the *continuation* epoch, coinciding with cessation of the first rhythm, and had a duration equal to 1 cycle of duple meter rhythm or 1 1/3 cycles of triple meter.

Contrast images (conditions 1 to 11 > rest) from the first level analyses were included in a second-level, random-effects analysis for group effects. Two behavioral covariates were also included in the second-level model: beat tapping consistency and BAT score. Beat tapping accuracy and GMSI scores were omitted as covariates as they were significantly correlated with beat tapping consistency and BAT scores, respectively. The two covariates included were the two (of four) that were least correlated with each other.

We had an *a priori* interest in striatal function, and preliminary results (e.g., whole brain contrasts for all rhythm-listening conditions vs. rest), which showed striatal activation was focused in the putamen (rather than caudate). Thus, region of interest (ROI) contrasts were completed for a bilateral putamen ROI (defined by the Automated Anatomical Labeling Toolbox, (Tzourio-Mazoyer et al., 2002) using a small-volume correction (SVC) with a voxel-level family-wise error (FWE) threshold of *p* < .05.

For contrasts between stages (e.g., *finding* > *continuation*, or *adjustment > rest*), individual conditions from simultaneous, staggered, congruent, and incongruent trials were weighted equally (for example, for *finding* > *continuation*, the three finding conditions were each weighted at 1/3, and the four finding conditions were each weighted at 1/4). For contrasts with respect to metrical incongruence, different stages were weighted equally across congruent and incongruent conditions (*fade-in* and *finding* epochs from staggered trials were omitted). For all contrasts, we used a cluster-level false discovery rate (FDR) threshold of *p* < .05, based on a cluster-forming threshold of < .0001 uncorrected (for individual stages > rest) or < .001 (for all other contrasts).

### Behavioral Analyses

For the beat-tapping task, we measured tapping consistency (coefficient of variation of inter-tap intervals) and tapping accuracy (mean absolute error between tap and beat times) for congruent and incongruent trials, as in previous studies (Cameron, Bentley, & Grahn, 2015; Cameron & Grahn, 2014). Mean ratings of beat strength for congruent and incongruent conditions were obtained for both pre-scan and during-scan rating sessions.

## Results

### Behavioral Results

Participants tapped the beat with greater accuracy (lower absolute tap-beat error) and greater consistency (lower CV of inter-tap intervals) for congruent compared to incongruent rhythms (accuracy: *t*(17) = 4.33, *p* < .001; consistency: *t*(17) = 7.72, *p* < .001). Tapping accuracy and consistency were correlated (Pearson’s *r* = .495, *p* = .018, 1-tailed). Congruent rhythms were rated as easier to maintain a sense of the beat, compared to incongruent rhythms, both before and during the scan (pre-scan: *t*(17) = 12.24, *p* < .001; in-scanner, *t*(10) = 4.51, *p* = .001, although in-scanner ratings from 7 participants were lost due to technical error). Neither measure of tapping performance correlated with either beat perception ability (BAT scores) or with musical training (GMSI scores). However, as expected, beat perception ability and musical training were correlated (Pearson’s *r* = .457, *p* = .028, 1-tailed). Behavioral results are shown in Figure 2.

**Figure 2.**
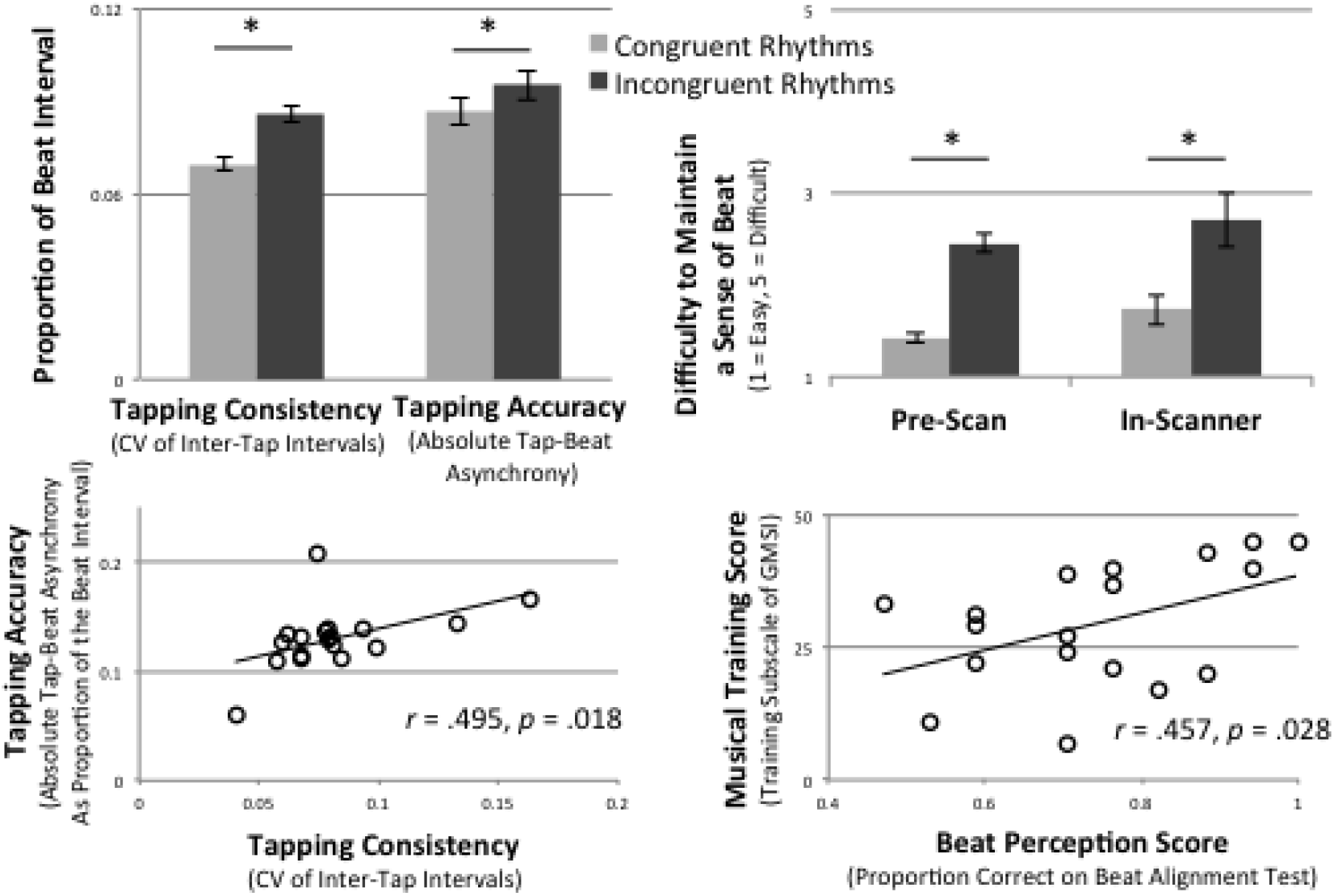
Behavioural results and correlations. Clockwise from top left: 1) Both tapping consistency (coefficient of variation, or CV, of inter-tap intervals) and tapping accuracy (absolute difference between tap and beat times as a proportion of the beat interval) were better for congruent compared to incongruent trials. 2) Congruent trials were also rated as have greater beat strength than incongruent trials (note that lower ratings reflect greater beat strength), for ratings made both before and during the fMRI scan (ratings did not differ between those two instances). Error bars indicate standard error of the mean, and * indicates *p* < .05. 3) Tapping accuracy and consistency were positively correlated. 4) Beat perception (proportion of correct trials on the BAT) and musical training (training subscale of the GMSI) were positively correlated.

### fMRI Results

Distinct regions of the striatum were differentially activated for the distinct stages of beat perception (see Table 1). During *beat finding*, when regularity was being detected, the dorsal putamen was more active than during *continuation*, when the beat had already been detected and was predictable. Although this activation was statistically significant for the right side, a subthreshold (SVC, FWE *p* = .076) peak was found for this contrast in the left dorsal putamen. In contrast, during *continuation*, the ventral putamen was more active (bilaterally) than during *finding*. In addition, this ventral region of the putamen was also more active for *beat adjustment* than both *continuation* (left) and *finding* (bilaterally). See Figure 3 (right panel) for images of activations from the putamen ROI contrasts (contrasts vs. rest are presented for clarity of visualization; peak voxels are similar to direct contrasts between the stages).

**Table 1.** Putamen Peaks Across Different Stages of Beat Perception.

| **Table 1.** |  |  |  |  |  |
| --- | --- | --- | --- | --- | --- |
| **Putamen Peaks Across Different Stages of Beat Perception** |  |  |  |  |  |
| **Finding > Continuation** |  |  |  |  |  |
| *Region of Putamen* | **x** | **y** | **z** | **t** | **p FWE** |
| R Dorsal | 24 | -1 | 13 | 3.92 | 0.013 |
| R Dorsal | 24 | 8 | 10 | 3.69 | 0.028 |
| **Continuation > Finding** |  |  |  |  |  |
| *Region of Putamen* | **x** | **y** | **z** | **t** | **p FWE** |
| L Ventral | -30 | -10 | -2 | 4.47 | 0.002 |
| L Ventral | -24 | 14 | -8 | 4.32 | 0.003 |
| R Ventral | 33 | 14 | -5 | 4.4 | 0.002 |
| R Ventral | 36 | -13 | -8 | 4.05 | 0.008 |
| **Adjustment > Finding** |  |  |  |  |  |
| *Region of Putamen* | **x** | **y** | **z** | **t** | **p FWE** |
| L Ventral | -30 | -10 | -5 | 6.6 | <.001 |
| L Ventral | -24 | 14 | -8 | 4.7 | <.001 |
| R Ventral | 33 | 14 | -5 | 5.1 | <.001 |
| R Ventral | 36 | -16 | -8 | 3.93 | 0.01 |
| **Adjustment > Continuation** |  |  |  |  |  |
| *Region of Putamen* | **x** | **y** | **z** | **t** | **p FWE** |
| L Ventral | -30 | -10 | -5 | 5.03 | <.001 |
| L Ventral | 30 | -1 | -5 | 4.4 | 0.002 |

Stereotactic coordinates (x, y, z) of peak voxels are in Montreal Neurological Institute space (mm).

**Figure 3.**
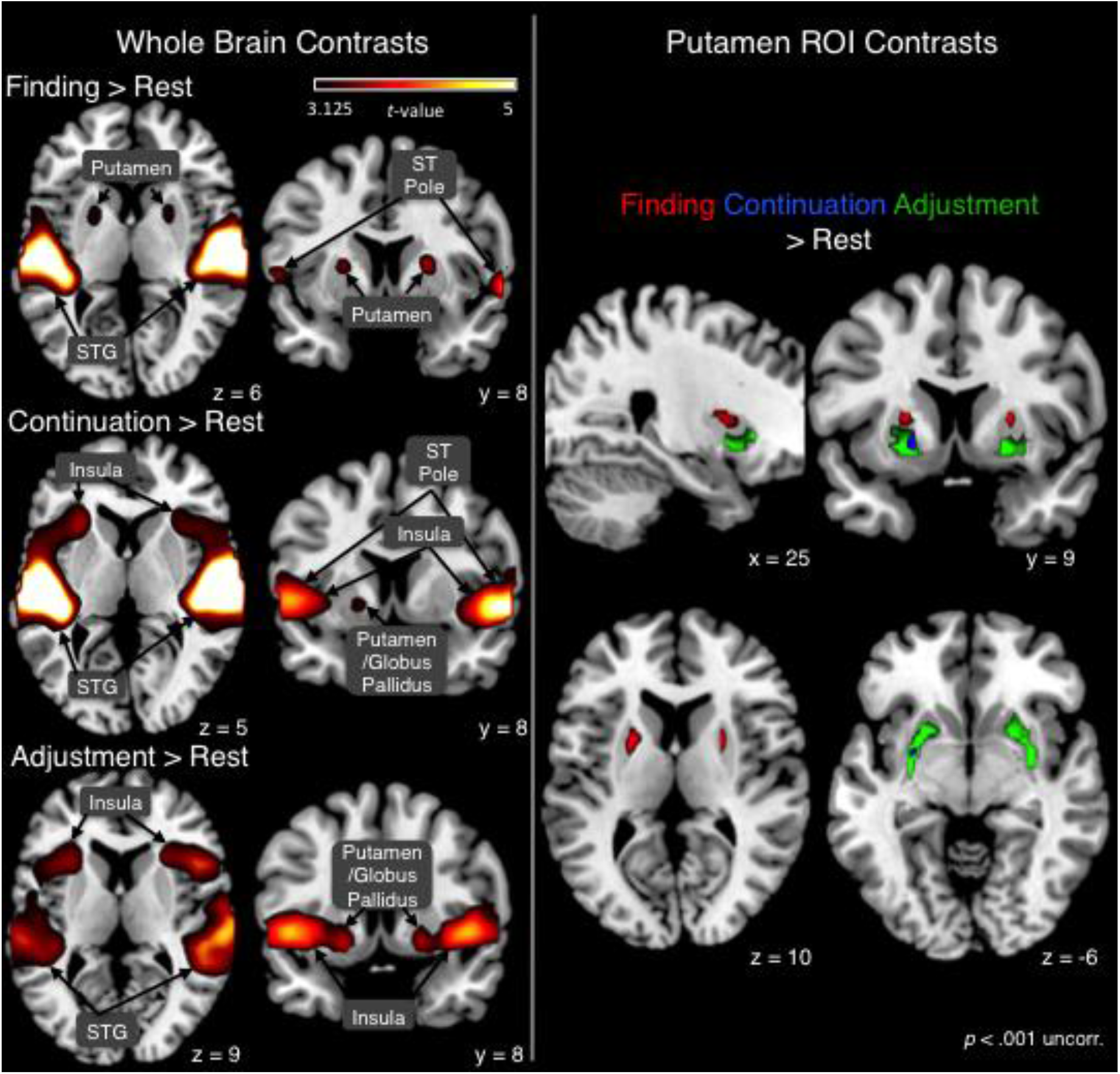
Contrasts for each stage (beat *finding*, beat *continuation*, and beat *adjustment*) vs. rest, for whole brain (left panel) and putamen ROI small-volume correction (right panel) analyses. Highlighted voxels are statistically significant, at *p* < .001 uncorrected. STG = superior temporal gyrus, ST Pole = superior temporal pole. See Table 2 for the complete list of results for whole brain analysis vs. rest.

At the whole-brain level, comparing the stages directly reveals regions that differ across stages (see Table 3 and Figure 4). For *beat finding* > *continuation*, the anterior and middle cingulate cortex had greater activity, as did left inferior parietal lobule (IPL), right supramarginal gyrus, cuneus and precuneus, and cerebellum (lobule 8). During *beat continuation* > *finding*, activity was greater in the insula, putamen, hippocampus, thalamus, inferior orbitofrontal cortex, and cerebellum (crus I). Regions with greater activity during *beat adjustment* > *finding* include insula, putamen, globus pallidus, hippocampus, and thalamus. Compared to *beat continuation, adjustment* was associated with greater activity in supplementary motor area (SMA), putamen, and right frontal operculum.

**Table 2.**
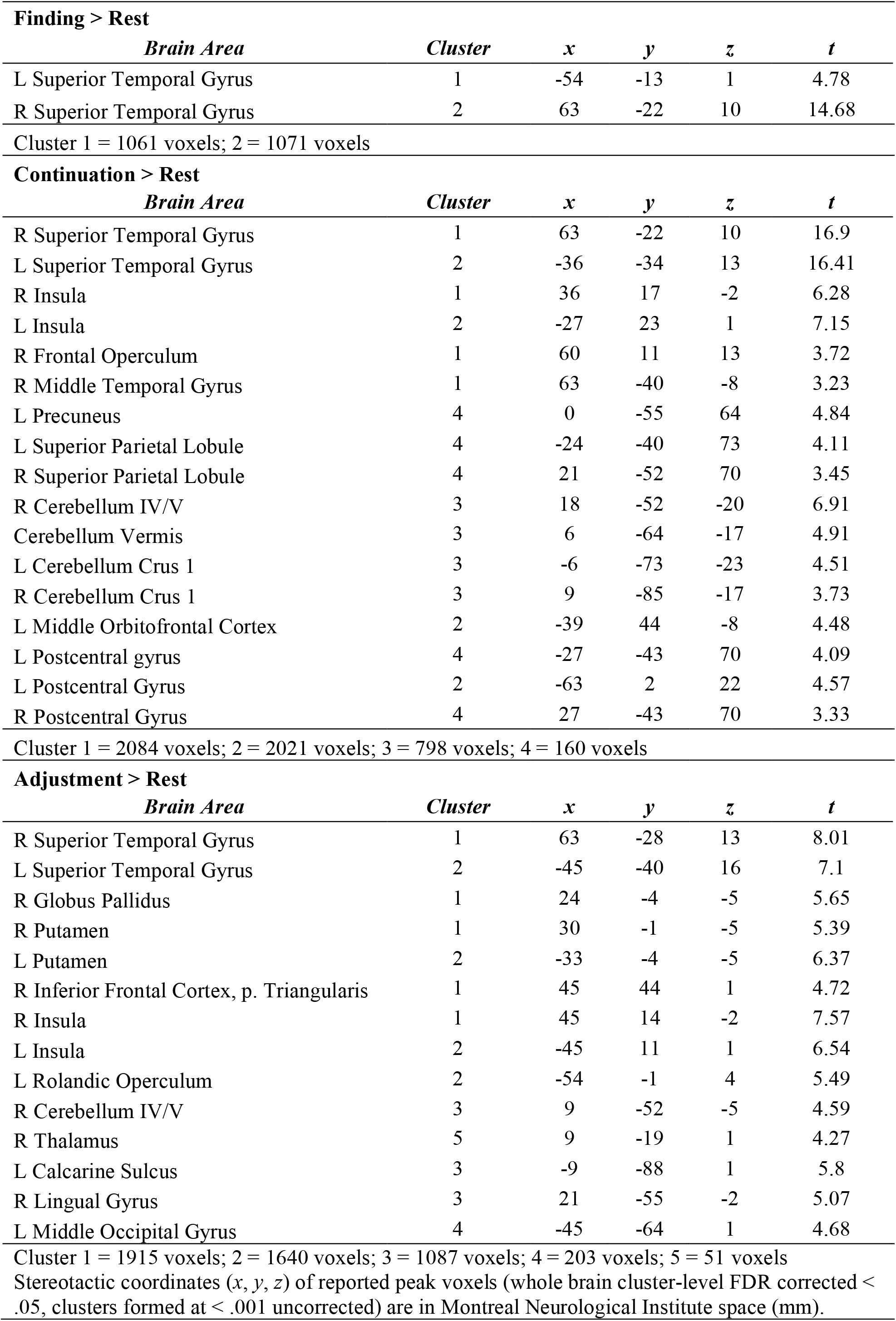
Peaks Across Different Stages of Beat Perception > Rest.

**Finding > Rest**
| *Brain Area* | *Cluster* | *x* | *y* | *z* | *t* |
| --- | --- | --- | --- | --- | --- |
| L Superior Temporal Gyrus | 1 | -54 | -13 | 1 | 4.78 |
| R Superior Temporal Gyrus | 2 | 63 | -22 | 10 | 14.68 |

**Continuation > Rest**
| *Brain Area* | *Cluster* | *x* | *y* | *z* | *t* |
| --- | --- | --- | --- | --- | --- |
| R Superior Temporal Gyrus | 1 | 63 | -22 | 10 | 16.9 |
| L Superior Temporal Gyrus | 2 | -36 | -34 | 13 | 16.41 |
| R Insula | 1 | 36 | 17 | -2 | 6.28 |
| L Insula | 2 | -27 | 23 | 1 | 7.15 |
| R Frontal Operculum | 1 | 60 | 11 | 13 | 3.72 |
| R Middle Temporal Gyrus | 1 | 63 | -40 | -8 | 3.23 |
| L Precuneus | 4 | 0 | -55 | 64 | 4.84 |
| L Superior Parietal Lobule | 4 | -24 | -40 | 73 | 4.11 |
| R Superior Parietal Lobule | 4 | 21 | -52 | 70 | 3.45 |
| R Cerebellum IV/V | 3 | 18 | -52 | -20 | 6.91 |
| Cerebellum Vermis | 3 | 6 | -64 | -17 | 4.91 |
| L Cerebellum Crus 1 | 3 | -6 | -73 | -23 | 4.51 |
| R Cerebellum Crus 1 | 3 | 9 | -85 | -17 | 3.73 |
| L Middle Orbitofrontal Cortex | 2 | -39 | 44 | -8 | 4.48 |
| L Postcentral gyrus | 4 | -27 | -43 | 70 | 4.09 |
| L Postcentral Gyrus | 2 | -63 | 2 | 22 | 4.57 |
| R Postcentral Gyrus | 4 | 27 | -43 | 70 | 3.33 |

**Adjustment > Rest**
| *Brain Area* | *Cluster* | *x* | *y* | *z* | *t* |
| --- | --- | --- | --- | --- | --- |
| R Superior Temporal Gyrus | 1 | 63 | -28 | 13 | 8.01 |
| L Superior Temporal Gyrus | 2 | -45 | -40 | 16 | 7.1 |
| R Globus Pallidus | 1 | 24 | -4 | -5 | 5.65 |
| R Putamen | 1 | 30 | -1 | -5 | 5.39 |
| L Putamen | 2 | -33 | -4 | -5 | 6.37 |
| R Inferior Frontal Cortex, p. Triangularis | 1 | 45 | 44 | 1 | 4.72 |
| R Insula | 1 | 45 | 14 | -2 | 7.57 |
| L Insula | 2 | -45 | 11 | 1 | 6.54 |
| L Rolandic Operculum | 2 | -54 | -1 | 4 | 5.49 |
| R Cerebellum IV/V | 3 | 9 | -52 | -5 | 4.59 |
| R Thalamus | 5 | 9 | -19 | 1 | 4.27 |
| L Calcarine Sulcus | 3 | -9 | -88 | 1 | 5.8 |
| R Lingual Gyrus | 3 | 21 | -55 | -2 | 5.07 |
| L Middle Occipital Gyrus | 4 | -45 | -64 | 1 | 4.68 |

Cluster 1 = 1915 voxels; 2 = 1640 voxels; 3 = 1087 voxels; 4 = 203 voxels; 5 = 51 voxels
Stereotactic coordinates (x, y, z) of reported peak voxels (whole brain cluster-level FDR corrected < .05, clusters formed at < .001 uncorrected) are in Montreal Neurological Institute space (mm).

**Table 3.**
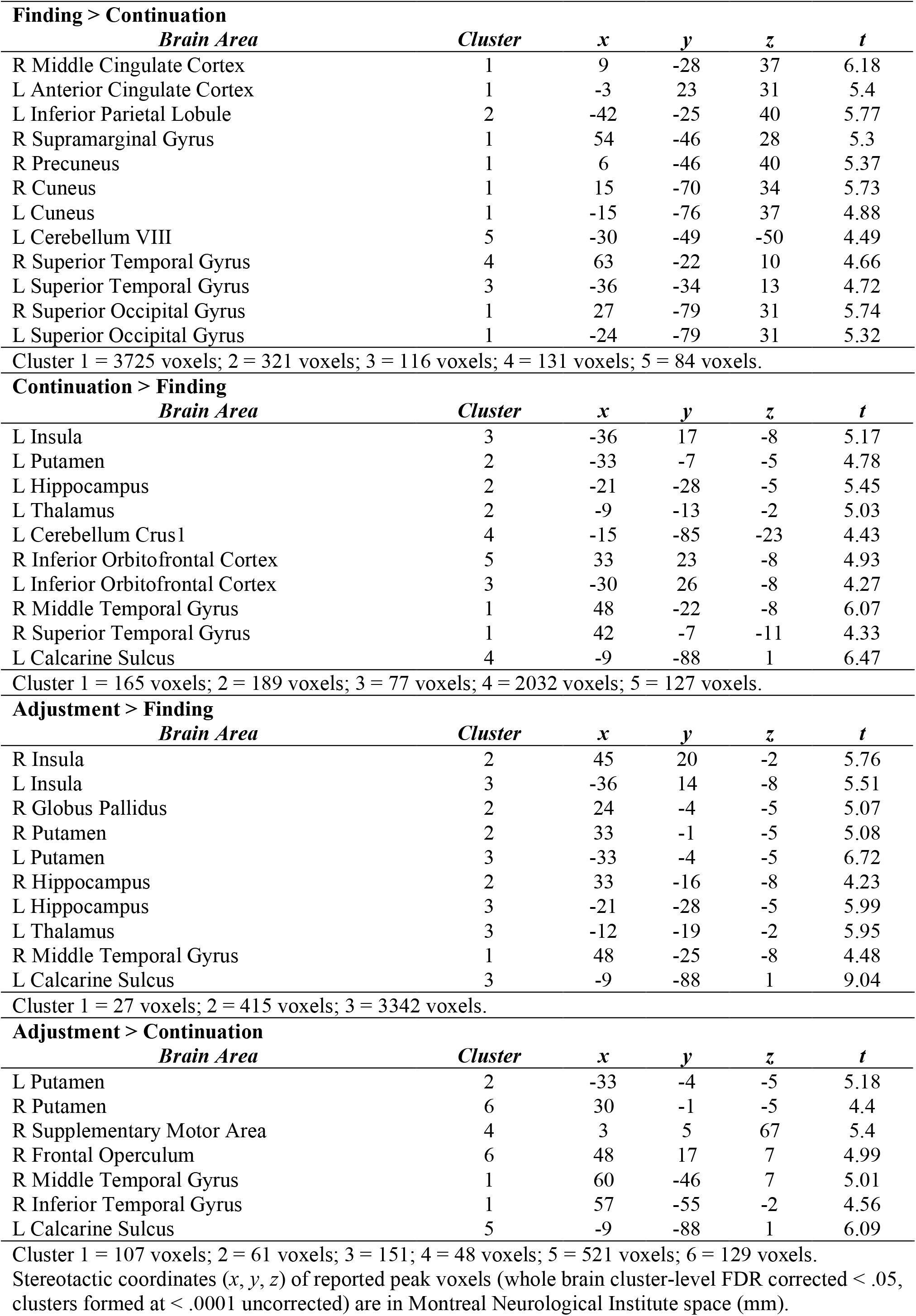
Peaks Across Different Stages of Beat Perception.

**Figure 4.**
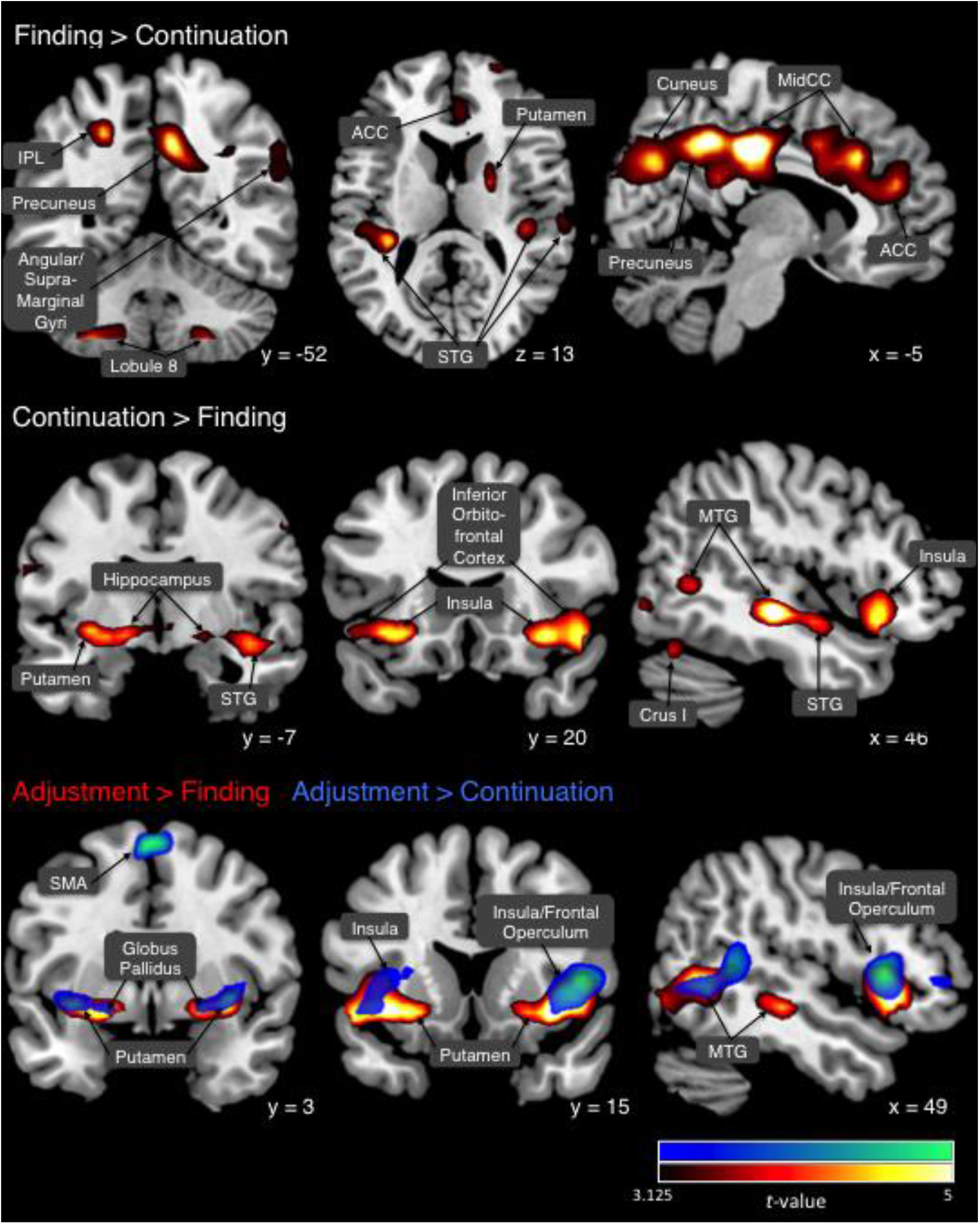
fMRI results from direct contrasts of stages of beat perception. Highlighted voxels are statistically significant, at *p* < .001 uncorrected.

Contrasting incongruent > congruent conditions revealed brain areas that were more active during listening to metrically incongruent (when multiple beat percepts were possible, corresponding to the differing metrical structures of the two stimulus rhythms) compared to metrically congruent rhythm pairs. Metrical incongruence was associated with activity in bilateral auditory and inferior frontal regions. During trials in which the two rhythms were metrically incongruent, compared to congruent trials, activity was greater in bilateral STG, left middle temporal gyrus (MTG), and right anterior insula as well as the frontal operculum (see Table 4 and Figure 5). The pattern of results (STG and insula/operculum) did not differ substantially for incongruent > congruent contrasts across the different stages (e.g., incongruent *adjustment* > congruent *adjustment*, and incongruent *finding* > congruent *finding*), although there was a subthreshold difference (incongruent > congruent) during *adjustment* in insula, and a significant effect during *finding* and *continuation* in STG (see Figure 5).

**Table 4.** Peaks During Incongruent > Congruent Rhythms (All Stages)

| *Brain Area* | *Cluster* | *x* | *y* | *z* | *t* |
| --- | --- | --- | --- | --- | --- |
| R Insula | 1 | 33 | 20 | 7 | 4.37 |
| R Frontal Operculum | 1 | 45 | 14 | 10 | 4.5 |
| R Superior Temporal Gyrus | 1 | 60 | -19 | 1 | 4.69 |
| L Superior Temporal Gyrus | 2 | -57 | -13 | 1 | 3.92 |
| L Middle Temporal Gyrus | 2 | -57 | -19 | 1 | 3.89 |

Cluster 1 = 505 voxels; 2 = 120 voxels.
Stereotactic coordinates (*x*, *y*, *z*) of reported peak voxels (whole brain cluster-level FDR corrected < .05, clusters formed at < .001 uncorrected) are in Montreal Neurological Institute space (mm).

**Figure 5.**
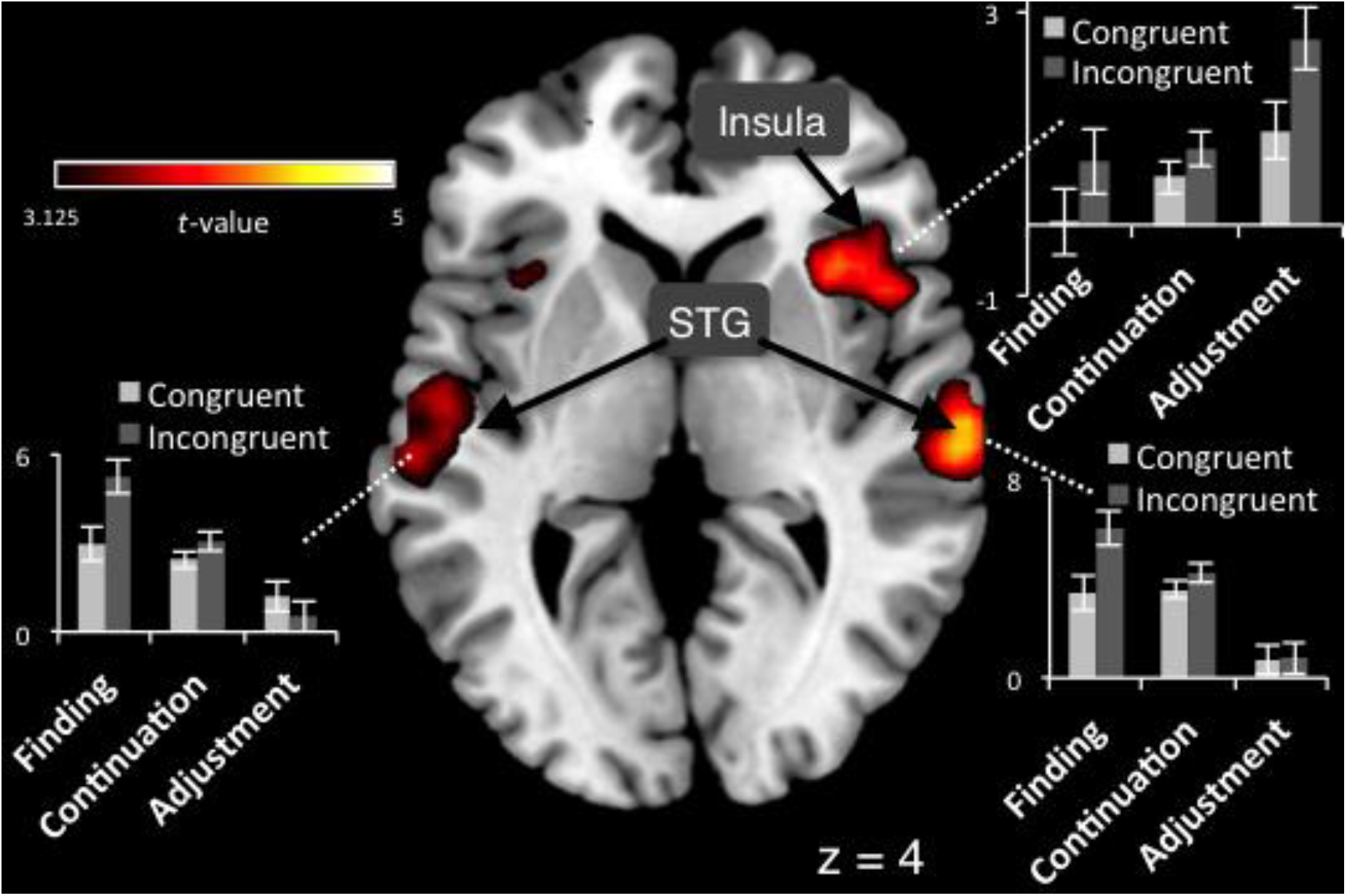
fMRI results contrasting listening during metrically incongruent > congruent trials. Highlighted voxels are statistically significant, at *p* < .001 uncorrected. Bar graphs indicate means and standard errors for incongruent > congruent contrasts for each individual stage (*beat finding, continuation, adjustment*), for peak voxels in the three regions.

Better beat tapping consistency and beat perception ability were associated with greater activity in temporal and parietal regions (see Table 5 and Figure 6). Activations in bilateral STG, right superior temporal pole, and left supramarginal gyrus were greater for participants with better beat tapping consistency (lower coefficient of variation of inter-tap intervals). Activations in right posterior STG and MTG were greater for participants with better beat perception (higher BAT scores).

**Table 5.** Correlations Between Behaviour and fMRI Activity.

| Table 5 |  |  |  |  |  |
| --- | --- | --- | --- | --- | --- |
| Correlations Between Behaviour and fMRI Activity |  |  |  |  |  |
| Tapping Consistency (CV) |  |  |  |  |  |
| *Brain Area* | *Cluster* | *x* | *y* | *z* | *t* |
| L Supramarginal gyrus | 1 | -57 | -25 | 19 | 3.96 |
| L Superior Temporal Gyrus | 1 | -54 | -13 | 1 | 4.78 |
| R Superior Temporal Gyrus | 2 | 63 | 2 | -5 | 4.36 |
| R Superior Temporal Pole | 2 | 60 | 8 | -8 | 4.14 |
| Cluster 1 = 97 voxels; 2 = 89 voxels. |  |  |  |  |  |
| Stereotactic coordinates (*x*, *y*, *z*) of reported peak voxels (whole brain cluster-level FDR corrected < .05, clusters formed at <.001 uncorrected) are in Montreal Neurological Institute space (mm). |  |  |  |  |  |
| BAT % correct |  |  |  |  |  |
| *Brain Area* | *Cluster* | *x* | *y* | *z* | *t* |
| R Superior Temporal Gyrus | 1 | 45 | -34 | 13 | 5.43 |
| R Middle Temporal Gyrus | 1 | 48 | -46 | 7 | 3.85 |
| Cluster 1 = 112 voxels. |  |  |  |  |  |

**Figure 6:**
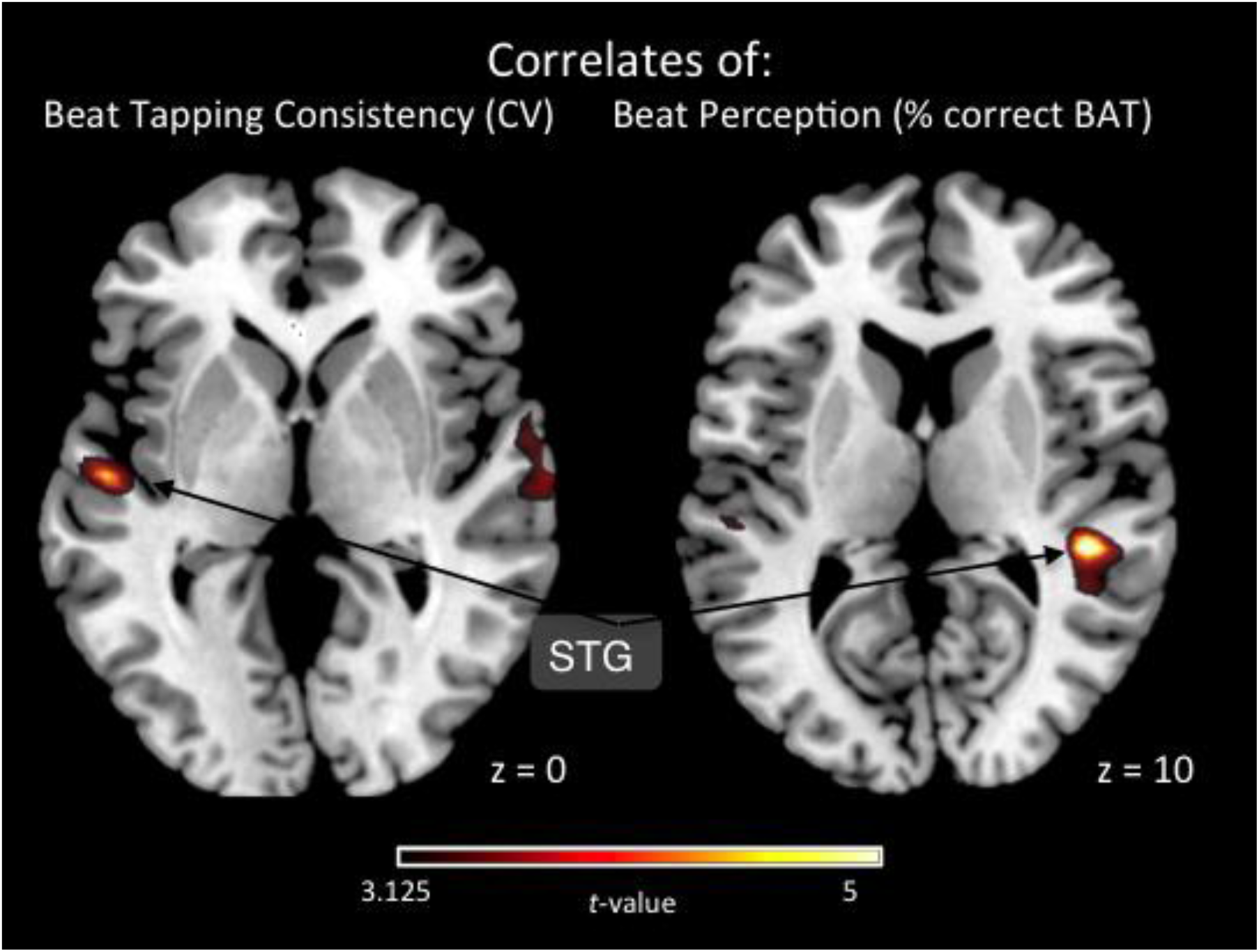
fMRI correlates of beat perception (% correct on BAT) and negative correlates beat tapping variability (CV of inter-tap intervals from beat tapping task). Highlighted voxels are statistically significant at *p* < .001 uncorrected.

As expected, comparison between tasks revealed greater activity in occipital cortex for the rating task compared to deviant-detection task (cluster-level FDR < .05).

## Discussion

### Differing neural mechanisms across stages of beat perception

The results show that distinct regions of the striatum respond during the different stages of beat perception, suggesting that distinct striatal functions occur at each stage. The dorsal putamen was most active during beat *finding*, which requires the detection of regularity. We propose that detection of regularity is supported by medium spiny neurons in the dorsal putamen that detect the coincidence of cortical oscillations (Matell & Meck, 2004). It may be that regularly-occurring coincidences between extrinsic and intrinsic oscillations initiate neural entrainment, in turn supporting the initiation of beat perception. The hemodynamic cost of neural populations in the dorsal putamen detecting regular oscillatory coincidence and entraining with other neural systems may be sufficient to underlie the observed fMRI data.

The ventral putamen was more active during *continuation* and even more so during *adjustment*, compared to *finding*, suggesting that that region supports ongoing temporal prediction during beat perception (Kotz, et al., 2009), and the integration of the temporal prediction errors (McClure et al., 2003) that occur during *adjustment*, into the ongoing, perceived beat structure.

“Actor” and “critic” functions have been associated with dorsal and ventral regions of the striatum, respectively, and these may correspond to their differential activation over the stages of beat perception. The actor-critic model suggests that the actor (dorsal) region of the striatum uses temporal prediction errors to modify stimulus-response associations in order to select actions, whereas the critic (ventral) region uses temporal prediction errors to update successive predictions based on the state of external-internal environment dynamics (O’Doherty et al., 2004; O’Doherty, Hampton, & Kim, 2007). Taken together with our results, this view suggests that temporal predictions and processing of prediction errors are ongoing functions of the striatum throughout beat perception—albeit with different functions. The “critic” (ventral putamen) is most active during *beat adjustment*, the stage requiring the integration of temporal prediction errors into ongoing predictions in order for beat perception (the internal state) to persist in the face of changing auditory rhythm (the external state). The “actor” (dorsal putamen) is more active during *beat finding*, the stage requiring the initial assessment of the stimulus in order to select the appropriate response (e.g., a beat rate to entrain to). Beat and meter perception can be thought of as a motoric and attentional behavior (London, 2004), and in that light, the notion of beat *finding* as modifying stimulus-response associations is apt. Thus, beat *finding* and *adjustment* may resemble previously proposed “actor-critic” functions of dorsal and ventral striatal regions, respectively.

Beyond striatal functions, the results suggest mechanisms involving other regions across the stages of beat perception. During beat *finding*, before the beat is detected, temporal intervals may be processed as absolute durations, rather than as durations relative to the beat—as no beat has yet been detected. Consistent with absolute timing occurring during *finding*, we found that activity in cerebellar lobule 8 was greater during *finding* than during *continuation*, as previous evidence shows that the cerebellum supports absolute timing (Teki et al., 2011). Additionally, during *finding*, attention orients in time to prospective beat positions. We found that activity in left inferior parietal cortex was greater during *finding* than *continuation*, consistent with previous evidence of its role in orienting attention to time (Bolger, Coull, & Schön, 2014; Coull & Nobre, 1998; Davranche, Nazarian, Vidal, & Coull, 2011; Cotti, Rohenkohl, Stokes, Nobre, & Coull, 2011). Moreover, during beat *finding*, the rhythmic stimulus is novel, and draws the listener’s attention. This general shift in attention to the rhythm may correspond to the greater anterior cingulate cortex activity in *finding* than *continuation*), given its role in attention to relevant external stimuli (Botvinick, Cohen, & Carter, 2004; Totah, Kim, Homayoun, & Moghaddam, 2009). Thus, the cerebellum, left inferior parietal cortex, and anterior cingulate may support the absolute timing, and temporal orienting of attention, that occur during beat *finding*.

After *finding*, during which the beat is detected and encoded, *beat continuation* enables the ongoing beat to be maintained and retrieved, in order to generate temporal predictions. The duration of the inter-beat interval is used to predict the onsets that will occur in the ongoing stimulus. The observed greater activation in the posterior hippocampus for *continuation* than *finding* may indicate support of the maintenance and retrieval of the beat interval, consistent with recent evidence of the posterior hippocampus’ role in auditory working memory, specifically the “analysis of auditory stimuli in real time” (Kumar et al., 2016). Additionally, previous research found hippocampal activation, though more anteriorly, during *beat continuation* (Grahn & Rowe, 2013).

The anterior insula is more active during both *continuation* and *adjustment* compared to *finding*, suggesting it supports ongoing beat perception. Insula activity has been widely observed in temporal auditory processing (Pastor, Macaluso, Day, & Frackowiak, 2006; Steinbrink, et al., 2009; Bamiou, et al., 2003), and during beat perception it may support the integration of auditory and motor processing (Kurth, Zilles, Fox, Laird, & Eickhoff, 2010; Mutschler et al., 2007; Mutschler et al., 2009; Zarate & Zatorre, 2005). The integration of auditory processing (i.e., rhythm perception) and motor processing (i.e., motoric entrainment, motor system activation) is the phenomenological hallmark of beat perception. However, auditory-motor integration is less involved in beat *finding*, during which auditory rhythm perception has not yet induced motoric entrainment, compared to *continuation* and *adjustment*. Thus, the observed anterior insula activation during *continuation* and *adjustment* (compared to *finding*) may be due to its role in auditory-motor integration.

### Metrical Incongruence

Human beat perception tends to persist during listening to metrically incongruent (e.g., polyrhythmic) stimuli. When listening to rhythms specifically designed to have an ambiguous metrical structure, allowing for multiple regularities to be perceived as the ‘beat’, listeners only track one rate as the ‘beat’ (Poudrier & Repp, 2013). This persistence of beat perception occurs in musical contexts in which different instruments produce distinct rhythms conforming to different metrical structures (metrical ambiguity, see London, 2004). When tapping the beat in metrically ambiguous compared to unambiguous rhythmic contexts, greater activity is found in inferior frontal regions, right anterior insula, and right inferior parietal cortex (Vuust, Roepstorff, Wallentin, Mouridsen, & Østergaard, 2006; Vuust, Wallentin, Mouridsen, Ostergaard, & Roepstorff, 2011), suggesting that these regions are part of a network relevant in integrating and cohering temporal information., Thus, we used metrically congruent and incongruent trials to compare the neural mechanisms of the maintenance of beat perception in the face of metrical incongruence in the rhythmic stimulus.

In addition to its differential activation across stages (greater in *continuation* and *adjustment* compared to *finding*), the right anterior insula was more active during metrically incongruent (vs. congruent) rhythms, when beat perception was persisting despite conflicting cues about beat structure. This is consistent with previous work on beat perception in metrically incongruent contexts, which found greater right anterior insula activity for incongruent compared to congruent contexts (Vuust et al., 2006). The function of the anterior insula in metrical incongruence may be similar to its function integrating auditory and motor processing in *continuation* and *adjustment* as discussed above. That is, metrical incongruence may require the anterior insula’s integration of auditory and motor processing to maintain beat perception, because of the difficulty in maintaining beat perception when there are multiple, conflicting metrical cues.

### Behavioral correlations

Correlations between behavioral and neural data showed that the STG was more active (bilaterally) for participants with superior beat tapping consistency (lower variability of inter-tap intervals) and (on the right side) for those with superior beat perception (greater proportion of correct trials in the BAT). Greater STG activity may be associated with more attention to the rhythms (Chapin et al., 2010), which likely supports performance on these tasks. In previous studies, STG activity correlated with metrical salience during synchronized tapping (Chen, Zatorre, & Penhune, 2006), with the degree of prediction in rhythm-synchronized tapping (Pecenka, Engel, & Keller, 2013), and with musical training (Koelsch, Fritz, Schulze, Alsop, & Schlaug, 2005). Musical training, and metrical salience and predictability of rhythms all likely support behavioural performance (musical training correlated with beat perception accuracy, see Supplemental Material), and may underlie behaviour-STG correlations.

## Conclusion

The striatum supports the processing of temporal regularity and the prediction of regular events. The results here indicate that dorsal and ventral regions of the putamen have distinct functions across the stages of beat perception. Dorsal putamen was more active for *beat finding* (possibly from detecting coincidence between, and phase resetting of, neural oscillations) and ventral putamen was more active during *beat adjustment* (possibly from processing of temporal prediction errors). Overall, we can place these striatal functions within the broader mechanisms that support the distinct processes of beat perception: mechanisms that vary in time (e.g., initial *beat finding* followed by *beat continuation*), mechanisms that depend on the unfolding and dynamic nature of the auditory rhythm (e.g., *beat adjustment*), and mechanisms that are arise from stimulus characteristics (e.g., metrical incongruence).

## Acknowledgements

This work was supported by funding from the Natural Sciences and Engineering Research Council of Canada

## Supplementary material

### Behavioural results

**Figure S1.**
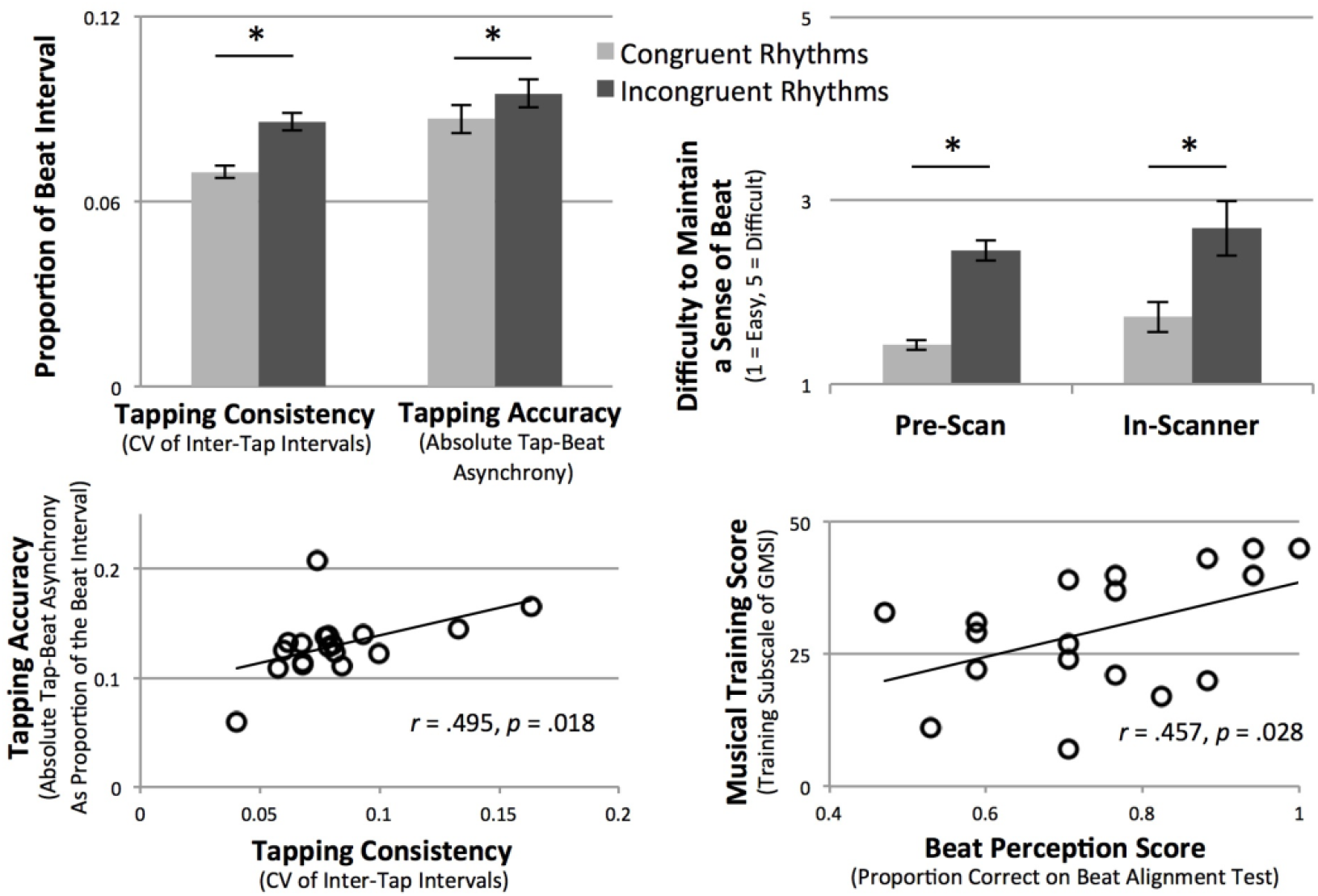
Behavioural results and correlations. Clockwise from top left: 1) Both tapping consistency (coefficient of variation, or CV, of inter-tap intervals) and tapping accuracy (absolute difference between tap and beat times as a proportion of the beat interval) were better for congruent compared to incongruent trials. 2) Congruent trials were also rated as have greater beat strength than incongruent trials (note that lower ratings reflect greater beat strength), for ratings made both before and during the fMRI scan (ratings did not differ between those two instances). Error bars indicate standard error of the mean, and * indicates *p* < .05. 3) Tapping accuracy and consistency were positively correlated. 4) Beat perception (proportion of correct trials on the BAT) and musical training (training subscale of the GMSI) were positively correlated.

## References

Aschersleben, G. (2002). Temporal control of movements in sensorimotor synchronization. Brain Cogn, 48(1), 66–79. doi:10.1006/brcg.2001.1304

Bolger, D., Coull, J. T., & Schön, D. (2014). Metrical rhythm implicitly orients attention in time as indexed by improved target detection and left inferior parietal activation. J Cogn Neurosci, 26(3), 593–605. doi:10.1162/jocn_a_00511

Botvinick, M. M., Cohen, J. D., & Carter, C. S. (2004). Conflict monitoring and anterior cingulate cortex: an update. Trends Cogn Sci, 8(12), 539–546. doi:10.1016/j.tics.2004.10.003

Cameron, D. J., Bentley, J., & Grahn, J. A. (2015). Cross-cultural influences on rhythm processing: reproduction, discrimination, and beat tapping. Front Psychol, 6, 366. doi:10.3389/fpsyg.2015.00366

Cameron, D. J., & Grahn, J. A. (2014). Enhanced timing abilities in percussionists generalize to rhythms without a musical beat. Front Hum Neurosci, 8, 1003. doi:10.3389/fnhum.2014.01003

Chapin, H. L., Zanto, T., Jantzen, K. J., Kelso, S. J., Steinberg, F., & Large, E. W. (2010). Neural responses to complex auditory rhythms: the role of attending. Front Psychol, 1, 224. doi:10.3389/fpsyg.2010.00224

Chen, J. L., Zatorre, R. J., & Penhune, V. B. (2006). Interactions between auditory and dorsal premotor cortex during synchronization to musical rhythms. Neuroimage, 32(4), 1771–1781. doi:10.1016/j.neuroimage.2006.04.207

Cotti, J., Rohenkohl, G., Stokes, M., Nobre, A. C., & Coull, J. T. (2011). Functionally dissociating temporal and motor components of response preparation in left intraparietal sulcus. Neuroimage, 54(2), 1221–1230. doi:10.1016/j.neuroimage.2010.09.038

Coull, J. T., & Nobre, A. C. (1998). Where and when to pay attention: the neural systems for directing attention to spatial locations and to time intervals as revealed by both PET and fMRI. J Neurosci, 18(18), 7426–7435.

Davranche, K., Nazarian, B., Vidal, F., & Coull, J. (2011). Orienting attention in time activates left intraparietal sulcus for both perceptual and motor task goals. J Cogn Neurosci, 23(11), 3318–3330. doi:10.1162/jocn_a_00030

Grahn, J., & Brett, M. (2007). Rhythm and beat perception in motor areas of the brain. J Cogn Neurosci, 19(5), 893–906. doi:10.1162/jocn.2007.19.5.893

Grahn, J., & Brett, M. (2009). Impairment of beat-based rhythm discrimination in Parkinson’s disease. Cortex, 45(1), 54–61. doi:10.1016/j.cortex.2008.01.005

Grahn, J. A., & Rowe, J. B. (2009). Feeling the beat: premotor and striatal interactions in musicians and nonmusicians during beat perception. J Neurosci, 29(23), 7540–7548. doi:10.1523/JNEUROSCI.2018-08.2009

Grahn, J. A., & Rowe, J. B. (2013). Finding and feeling the musical beat: striatal dissociations between detection and prediction of regularity. Cereb Cortex, 23(4), 913–921. doi:10.1093/cercor/bhs083

Koelsch, S., Fritz, T., Schulze, K., Alsop, D., & Schlaug, G. (2005). Adults and children processing music: an fMRI study. Neuroimage, 25(4), 1068–1076. doi:10.1016/j.neuroimage.2004.12.050

Kotz, S. A., Schwartze, M., & Schmidt-Kassow, M. (2009). Non-motor basal ganglia functions: a review and proposal for a model of sensory predictability in auditory language perception. Cortex, 45(8), 982–990. doi:10.1016/j.cortex.2009.02.010

Kumar, S., Joseph, S., Gander, P. E., Barascud, N., Halpern, A. R., & Griffiths, T. D. (2016). A Brain System for Auditory Working Memory. J Neurosci, 36(16), 4492–4505. doi:10.1523/JNEUROSCI.4341-14.2016

Kung, S. J., Chen, J. L., Zatorre, R. J., & Penhune, V. B. (2013). Interacting cortical and basal ganglia networks underlying finding and tapping to the musical beat. J Cogn Neurosci, 25(3), 401–420. doi:10.1162/jocn_a_00325

Kurth, F., Zilles, K., Fox, P. T., Laird, A. R., & Eickhoff, S. B. (2010). A link between the systems: functional differentiation and integration within the human insula revealed by meta-analysis. Brain Struct Funct, 214(5-6), 519–534. doi:10.1007/s00429-010-0255-z

Large, E. W., & Kolen, J. F. (1994). Resonance and the perception of musical meter. Conn Sci, 6(2-3), 177–208.

Large, E., & Snyder, J. (2009). Pulse and meter as neural resonance. Ann N Y Acad Sci, 1169, 46–57. doi:NYAS04550

London, J. (2004). Hearing in time: Psychological aspects of musical meter. Oxford University Press.

Matell, M. S., & Meck, W. H. (2004). Cortico-striatal circuits and interval timing: coincidence detection of oscillatory processes. Brain Res Cogn Brain Res, 21(2), 139–170. doi:10.1016/j.cogbrainres.2004.06.012

McClure, S. M., Berns, G. S., & Montague, P. R. (2003). Temporal prediction errors in a passive learning task activate human striatum. Neuron, 38(2), 339–346.

Müllensiefen, D., Gingras, B., Musil, J., & Stewart, L. (2014). The musicality of non-musicians: an index for assessing musical sophistication in the general population. PLoS One, 9(2), e89642. doi:10.1371/journal.pone.0089642

Mutschler, I., Schulze-Bonhage, A., Glauche, V., Demandt, E., Speck, O., & Ball, T. (2007). A rapid sound-action association effect in human insular cortex. PLoS One, 2(2), e259. doi:10.1371/journal.pone.0000259

Mutschler, I., Wieckhorst, B., Kowalevski, S., Derix, J., Wentlandt, J., Schulze-Bonhage, A., & Ball, T. (2009). Functional organization of the human anterior insular cortex. Neurosci Lett, 457(2), 66–70. doi:10.1016/j.neulet.2009.03.101

O’Doherty, J., Dayan, P., Schultz, J., Deichmann, R., Friston, K., & Dolan, R. J. (2004). Dissociable roles of ventral and dorsal striatum in instrumental conditioning. Science, 304(5669), 452–454. doi:10.1126/science.1094285

O’Doherty, J. P., Hampton, A., & Kim, H. (2007). Model-based fMRI and its application to reward learning and decision making. Ann N Y Acad Sci, 1104, 35–53. doi:10.1196/annals.1390.022

Pastor, M. A., Macaluso, E., Day, B. L., & Frackowiak, R. S. (2006). The neural basis of temporal auditory discrimination. Neuroimage, 30(2), 512–520. doi:10.1016/j.neuroimage.2005.09.053

Pecenka, N., Engel, A., & Keller, P. E. (2013). Neural correlates of auditory temporal predictions during sensorimotor synchronization. Front Hum Neurosci, 7, 380. doi:10.3389/fnhum.2013.00380

Poudrier, E., & Repp, B. H. (2013). Can musicians track two different beats simultaneously? Music Perception: An Interdisciplinary Journal, 30(4), 369–390.

Tal, I., Large, E. W., Rabinovitch, E., Wei, Y., Schroeder, C. E., Poeppel, D., & Golumbic, E. Z. (2017). Neural Entrainment to the Beat: the “Missing Pulse” Phenomenon. J Neurosci, 37(26), 6331–6341.

Teki, S., Grube, M., Kumar, S., & Griffiths, T. D. (2011). Distinct neural substrates of duration-based and beat-based auditory timing. J Neurosci, 31(10), 3805–3812. doi:10.1523/JNEUROSCI.5561-10.2011

Totah, N. K., Kim, Y. B., Homayoun, H., & Moghaddam, B. (2009). Anterior cingulate neurons represent errors and preparatory attention within the same behavioral sequence. J Neurosci, 29(20), 6418–6426. doi:10.1523/JNEUROSCI.1142-09.2009

Tzourio-Mazoyer, N., Landeau, B., Papathanassiou, D., Crivello, F., Etard, O., Delcroix, N., … Joliot, M. (2002). Automated anatomical labeling of activations in SPM using a macroscopic anatomical parcellation of the MNI MRI single-subject brain. Neuroimage, 15(1), 273–289. doi:10.1006/nimg.2001.0978

Vuust, P., Roepstorff, A., Wallentin, M., Mouridsen, K., & Østergaard, L. (2006). It don’t mean a thing… Keeping the rhythm during polyrhythmic tension, activates language areas (BA47). Neuroimage, 31(2), 832–841. doi:S1053-8119(05)02568

Vuust, P., Wallentin, M., Mouridsen, K., Ostergaard, L., & Roepstorff, A. (2011). Tapping polyrhythms in music activates language areas. Neurosci Lett, 494(3), 211–216. doi:10.1016/j.neulet.2011.03.015

Zarate, J. M., & Zatorre, R. J. (2005). Neural substrates governing audiovocal integration for vocal pitch regulation in singing. Ann N Y Acad Sci, 1060, 404–408. doi:10.1196/annals.1360.058

Zarco, W., Merchant, H., Prado, L., & Mendez, J. C. (2009). Subsecond timing in primates: comparison of interval production between human subjects and rhesus monkeys. J Neurophysiol, 102(6), 3191–3202. doi:10.1152/jn.00066.2009

